# Theory of Mind*s*: Managing mental state inferences in working memory is associated with the dorsomedial subsystem of the default network and social integration

**DOI:** 10.1101/697391

**Authors:** Meghan L. Meyer, Eleanor Collier

## Abstract

We often interact with multiple people at a time and consider their various points-of-view to facilitate smooth social interaction. Yet, how our brains track multiple mental states at once, and whether skill in this domain links to navigating real-world social interactions, remains underspecified. To fill this gap, we developed a novel social working memory paradigm in which participants manage two- or four-people’s mental states in working memory, as well as control trials in which they alphabetize two- or four-people’s names in working memory. In Study 1, we found that the dorsomedial subsystem of the default network shows relative increases in activity with more mental states managed in working memory. In contrast, this subsystem shows relative decreases in activity with more non-mental state information (the number of names alphabetized) managed in working memory. In Study 2, only individual differences in managing mental states in working memory, specifically on trials that posed the greatest mental state load to working memory, correlated with social network integration. Collectively, these findings add further support to the hypothesis that social working memory relies on partially distinct brain systems and may be a key ingredient to success in a social world.

---

Imagine the following scenarios: a business meeting in which employees and bosses keep track of multiple colleagues’ intentions; an interracial interaction in which individuals manage their stereotypes and anxieties during social inference; a gossip session in which friends tailor what is said based on one another’s knowledge of the situation. These scenarios are tied by a common theme: people actively manage a great deal of social information in mind to ensure smooth social interactions. How do we pull off such complex social information processing on the fly?

The maintenance and manipulation of high-level social information (e.g., personality traits, mental states, interpersonal relationships) has been characterized as social working memory (SWM) and there is evidence that this process relies on partially distinct brain systems from those supporting non-social forms of working memory. Specifically, regions of the brain’s default network, which includes medial prefrontal cortex (MPFC), dorsomedial prefrontal cortex (dMPFC), precuneus/posterior cingulate (PC/PCC), tempoparietal junction (TPJ), and temporal poles (TPs), systematically increase activation when participants maintain and manipulate high-level social information in their minds (i.e., information about friends’ personalities; Meyer, Spunt, Berkman, Taylor, & Lieberman, 2012; Meyer, Taylor, & Lieberman, 2015). In contrast, this same network is well-known to systematically disengage during non-social forms of working memory (Anticevic, Repovs, Shulman, & Barch, 2010; McKiernan, Kaufman, Kucera-Thompson, & Binder, 2003). In fact, the default network even shows this differential relationship when participants process the same stimuli either socially or non-socially in working memory. In one study, participants completed working memory trials in which they were shown two, three, or four of their own friends’ names and next either ranked them along trait dimensions (e.g., who is the funniest?) or alphabetized them (e.g., whose name is first alphabetically?). Within the same participants, the default network parametrically increased activation as a function of the number of friends’ traits considered in working memory even though it parametrically decreased activation as a function of the number of friends’ names alphabetized in working memory (Meyer et al., 2015). Thus, the default network appears to be differentially associated with high-level social and non-social information processing in working memory.

While these findings are provocative, they generate many more questions than answers. A particularly important question to answer is whether the results generalize to multiple types of high-level social information processing managed in working memory. In fact, at first blush, it appears as though the findings may not generalize to managing others’ mental states (as opposed to traits) in working memory. Specifically, maintaining others’ emotional facial expressions in working memory is associated with increased activity in the lateral frontoparietal network (Smith et al., 2017), but relatively suppressed activity in the default network (Xin & Lei, 2015). However, in these paradigms, participants are asked to judge whether a facial expression matches the facial expression shown in a previous trial. Thus, task performance does not require managing mental state *inferences* in working memory; rather, only externally presented facial features need to be considered. As a result, it remains unclear whether the default network may increase activity in response to managing mental state inferences in working memory.

Another important question to answer is whether SWM skills relate to social integration. It has been suggested that just as non-social working memory (non-SWM) skills relate to academic success (Conway, Conwan, Bunting, Therriault, & Minkoff, 2002; Rohde & Thompson, 2007; St. Clair-Thompson & Gathercole, 2006), SWM skills may relate to interpersonal success (Meyer et al., 2012; Meyer et al., 2015; Krol, Meyer, Lieberman, & Bartz, 2018; Smith, Killgore, & Alkozei, & Lane, 2018a; Mikels, Reuter-Lorenz, 2019). Indeed, individuals who can accurately manage more high-level social information in mind should make the most accurate predictions about others; and critically, feeling accurately understood by peers promotes friendship with them (Cross, Bacon, & Morris, 2000; Reis & Shaver, 1988). Yet, pinpointing whether SWM accuracy relates to social integration has been challenging, in part, because responses to the SWM paradigm previously used to assess the relationship with social integration are subjective. That is, past work has linked social integration to SWM performance on the trait ranking paradigm, in which participants rank their friends from ‘most-to-least’ along trait dimensions over a delay period (Krol et al., 2018). However, when a participant indicates that one of their friends is funnier than the other on a given SWM trial, it is unclear whether or not they answered accurately. What is needed is a SWM paradigm in which task accuracy can be computed objectively, so that individual differences in SWM capacity can be linked to differences in social integration.

The goal of the present research was to create a new SWM paradigm to fill these gaps. In this paradigm, participants first watch a video montage, which introduces a social network of characters with different interpersonal relationships (i.e., friends, lovers, competitors, and enemies). Participants next complete a SWM task in which they are asked to determine how a given character would feel, based on other characters’ feelings. Critically, the number of characters shown varies from trial-to-trial, with half of the trials showing two characters and the other half showing four characters. This allows for the isolation of neural activity and task performance as a function of SWM load, above and beyond mental state inference per se. Because interpersonal relationships tend to create known emotional contingencies between people (e.g., people tend to be sad to learn their friends are unhappy but often delight to hear their enemies are unhappy (Cikara, Botvinick, & Fiske, 2011)), we were able to measure SWM task accuracy more objectively. The paradigm also included non-SWM trials that were matched on a number of features (e.g., task difficulty), but that did not induce mental state inference. With this paradigm, we were able to test whether considering multiple mental state inferences in working memory differentially engages the default network (Study 1) and whether SWM task accuracy uniquely predicts social integration (Study 2).

## Study 1

### Participants

Thirty-seven participants (13 male, 23 female, 1 unspecified; Mean age=29.32 years, SD=11.32; 65% Caucasian, 19% Asian, 5% Hispanic or Latino, 11% other) completed Study 1 for course credit or monetary payment ($20/hour). Participants were eligible to participate if they did not have any metal in their body, were not claustrophobic, and were right-handed. Sample size was determined by available funding; and a priori power analysis in G*Power 3.1 (Faul, Erdfelder, Lang, & Buchner, 2007) showed that this sample size is estimated to provide 99% power to detect a medium within-subject working memory type x load level interaction effect (i.e., np2=.09). Participants provided informed consent in accordance with the Dartmouth College Institutional Review Board (IRB).

### Procedure

#### Social Working Memory Paradigm

Directly before fMRI scanning, participants first watched a montage introducing them to characters from the television show *Orange is the New Black*. A professional television editor developed the montage to help ensure that it effectively communicated the interpersonal relationships between the characters in an engaging format. Five characters were introduced and relationships between the characters included friends, enemies, lovers, and competitors. Before running the study, a sample of pilot subjects watched the montage and next described the relationships between the characters (N=19). All participants accurately described the relationships, suggesting that the relationship dynamics are well conveyed by the montage.

In Study 1, after watching the montage, participants indicated whether or not they had previously seen the television show *Orange is the New Black*. This allowed us to subsequently examine whether task performance varies as a function of familiarity with the characters in the montage. Participants next received instructions regarding how to complete the SWM and non-SWM trials and completed six, unique practice trials (2 per condition) on a laptop so that they were familiar with the task prior to scanning.

While undergoing fMRI scanning, participants completed the primary task (Figure 1), which employs a 2 (working memory type: SWM vs. non-SWM) x 2 (working memory load: two vs. four) experimental design. Participants completed 36 randomly presented trials (9 per condition). For two-load SWM trials, participants see two characters on the screen (4 seconds). The mental state of one character is provided, indicated by a thumbs-up (positive mental state) or thumbs-down (negative mental state). The mental state of the other ‘target’ character is not provided. Next, participants are instructed to consider how the target character would feel, based on the other character’s given mental state and relationship with the target character during a delay period (4 seconds). For example, if the two characters are friends and the character with the shown mental state is feeling positive, then the participant would reason that the other character, who likes their friend, would feel positive. After making the rating on a button box, the trial advanced to jittered fixation for a randomly generated duration between the range of 1.55-4.47 seconds (M=2.94 seconds).

**Figure 1.**
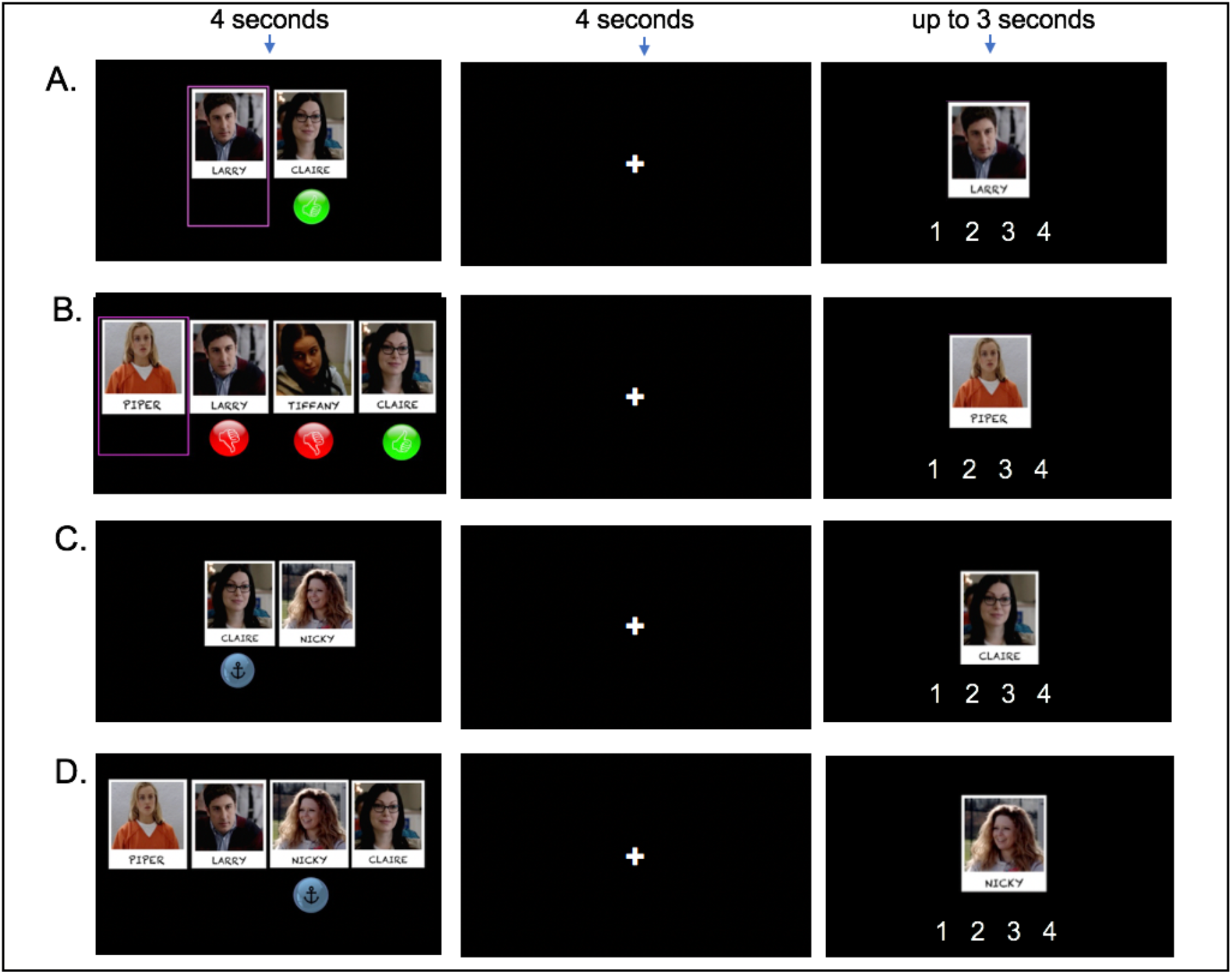
Social Working Memory (SWM) Paradigm. For SWM trials (Panels A-B), participants determine how the target character (pink box) would feel, based on the other characters’ feelings. The red thumbs-down sign indicates that a character is feeling negatively, whereas the green thumbs-up sign indicates that a character is feeling positively. For non-SWM trials (Panels C-D), participants alphabetize the characters’ names, based on whose name anchors the alphabet line (which is indicated by the anchor sign). Panels A and C show two-load working memory trials and Panels B and D show four-load working memory trials. Participants encode the initial stimuli for 4 seconds, which is followed by a 4 second delay period. Participants next have up to 3 seconds to make their response.

Four-load SWM trials have a similar format. The key difference is that for these trials participants are shown *three* characters with mental states and one target character with no mental state. During the subsequent delay period, participants reason how the target character would feel, based on the other characters’ feelings. Thus, two-load and four-load trials are identical, except that the four-load trials afford greater SWM load. To further help determine task accuracy objectively, participants were instructed to reason about the mental states serially and ‘from left to right.’ That is, in the example shown in Figure 1B, participants would reason about Piper’s mental state in the following order: Larry’s influence on her mental state (she would be unhappy if Larry, her fiancé, was unhappy; thus a rating of 1), her happiness would be taken up a notch if she find out that her enemy Tiffany was unhappy (now a rating of 2), and up another notch if she found out Claire, her girlfriend, is happy (now a rating of 3, the final, correct answer for this trial). In this way, mental state inferences proceed one at a time, and each additional character’s mental state is considered independently of one another.

Non-SWM trials have the same format, except here participants alphabetize the characters’ names over a delay period, also based on a (non-social) relationship between stimuli (i.e., the relationship between the position of letters on the alphabet line). Participants see a set of characters and an anchor sign is shown under one character (4 secs). Participants are told that the anchor indicates that for a given trial, the alphabet line should start with the first letter of the character’s name with the anchor. For example, if the anchor appeared under the character named Piper, ‘P’ should be treated as the first letter of the alphabet, and once reaching ‘Z’ the alphabet line would wrap around so that ‘O’ would become the last letter of the alphabet. During the delay period, participants alphabetize the characters’ names based on this relational rule between stimuli. To ensure that the answer could vary for two-load trials, participants were shown the character with the anchor for half of the probe-responses and the target character for the other half of the probe-responses. As is the case in the SWM trials, the four-character trials were identical to the two-character trials, but simply afforded a greater working memory load. In their probe response, participants indicated the alphabetical position of the shown character. This control condition was chosen to be consistent with past, non-social working memory research. Many working memory studies have parameterized the amount of alphabetizing over a delay period to assess the brain systems associated with working memory (e.g., D’Esposito, Postle, & Lease, 1999; Postle, Berger, & D’Esposito, 1999; Fougnie & Marois, 2007; Maniscalco & Lau, 2015; Postle, Ferrarelli, Hamidi, & Feredoes, 2006). Creating this alphabetizing condition allowed us to be consistent with prior research, while also having participants reason based on relationships between encoded stimuli.

#### fMRI Scanning

Scanning was conducted with a Siemens Trio 3T Prisma. Participants first completed a T2-weighted structural scan acquired coplanar with the functional images and with the following parameters: TR=2300 ms, TE=2.32 ms, 0.9 mm slice thickness, FOV=24 cm, matrix=256 × 256, flip angle=8°. During this time, participants watched the *Orange is the New Black* montage for a second time to refresh their memory of the relationships between characters. The task was completed during two functional runs using an EPI gradient-echo sequence with the following parameters: TR=1000 ms, TE=30 ms, 2.5 mm slice thickness, FOV=24 cm, matrix=96 × 96 and flip angle=59°.

#### fMRI Data Analysis

Functional brain imaging data was first reoriented in SPM8 (Wellcome Department of Cognitive Neurology, Institute for Neurology, London, United Kingdom) and skull-stripped with Brain Extraction Tool (BET; Smith, 2002) in FSL. All subsequent fMRI analyses were performed with SPM8. Preprocessing steps included spatial realignment to correct for head motion, normalization into a standard stereotactic space as defined by the Montreal Neurological Institute, and spatial smoothing using an 8-mm Gaussian kernel, full width at half-maximum.

For each subject, neural activity was modeled for each condition using a general linear model. Because we observed differences in reaction time (RT) across our SWM and non-SWM conditions (see Results section below), we also ran models that included, for each condition, RT as a parametric modulator. These models were run to ensure that clusters that varied between content (SWM vs. non-SWM) and load level (two-load vs. four-load) were not conflated with differences in RT across these conditions. Participants’ first-level models included a regressor for each condition of interest (convolved with the hemodynamic response function (HRF) vs. implicit baseline (i.e. SWM two-load vs. implicit baseline; SWM four-load vs. implicit baseline; non-SWM two-load vs. implicit baseline; non-SWM four-load vs. implicit baseline), as well as six motion regressors for each of the motion parameters from image realignment.

#### Whole-Brain Analyses

Next, we used a flexible factorial design to identify clusters of neural activity across subjects that are differentially associated with four-load (vs. two-load) SWM trials relative to four-load (vs. two-load) non-SWM trials. For completion, we also tested the effect of working memory content (SWM vs. non-SWM) and load level (two characters vs. four characters). Whole-brain results were voxelwise thresholded at p<.001, with a cluster-extent threshold corrected for family wise error rate (FWE), using a cluster-defining FWE threshold of p<.001, as determined by SPM. This approach corresponded with a cluster-extent threshold of 352 voxels.

#### Pre-defined Default Network Analyses

We followed-up our whole-brain interaction results with analyses performed with pre-defined default network subsystems. We used the default network subsystems defined by Yeo et al. (2011), which were generated from 1,000 participants’ resting state scans, and are thus highly reliable. This approach allowed us to assess whether 1) our results are replicated in a formally defined default network and 2) the interaction we observed in the dorsomedial subsystem in the whole-brain analysis was indeed reflected by relatively greater activity to SWM four-load (vs. two-load) trials and relatively less activity to non-SWM four-load (vs. two-load) trials, using clusters defined independently of the whole-brain results.

We extracted parameter estimates from each condition (vs. implicit baseline) for each of the three default network subsystems identified by Yeo et al. (2011). The subsystems include the dorsomedial subsystem (dorsomedial prefrontal cortex (DMPFC), temporoparietal junction (TPJ), middle temporal gyrus (MTG) extending into temporal pole (TP), and inferior frontal gyrus (IFG)), core subsystem (medial prefrontal cortex (MPFC), posterior cingulate/precuneus (PC/PCC), posterior inferior parietal lobule (PIPL)), and medial temporal lobe subsystem (hippocampal formation, retrosplenial cortex, and dorsal posterior inferior parietal lobule (dPIPL); Figure 4A). We then tested the interaction and appropriate follow-up t-tests for each network subsystem, separately. Supplementary Figure 1 shows the overlap between the clusters observed in the whole-brain interaction and those identified by Yeo et al. (2011) as part of the dorsomedial subsystem, which includes substantial overlap in DMPFC, TPJ, and temporal poles.

### Results

#### Task Performance

Task accuracy was defined as the percentage of correctly answered trials for each condition. There was no significant interaction between working memory content (SWM vs. non-SWM) and load level (two-load vs. four-load; *F*(1,36)=.05, *p*=.820, np2=.001; Figure 2A), suggesting that task performance was equivalent between the SWM and non-SWM trials. Consistent with this, there was no main effect of working memory content (SWM vs. non-SWM; *F*(1,36)=2.97, *p*=.094, np2=.08). In contrast, there was a significant main effect of load level (*F*(1,36)=81.90, *p*<.0001, np2=.70) such that four-load trials were more challenging (M=55%, SD=24%) than two-load trials (M=82%, SD=17%). Task accuracy did not statistically differ between participants who had previously seen *Orange is the New Black* (N=15) relative to participants who had not seen the show (N=22; *p*’s across load level comparisons>.177, Cohen’s d’s<.42; See *Supplementary Table 1*). Collectively, these results suggest that task accuracy was equivalent for the SWM and non-SWM trials, that the four-load manipulation successfully taxes SWM and non-SWM, and that prior exposure to the television show *Orange is the New Black* does not moderate task performance.

**Figure 2.**
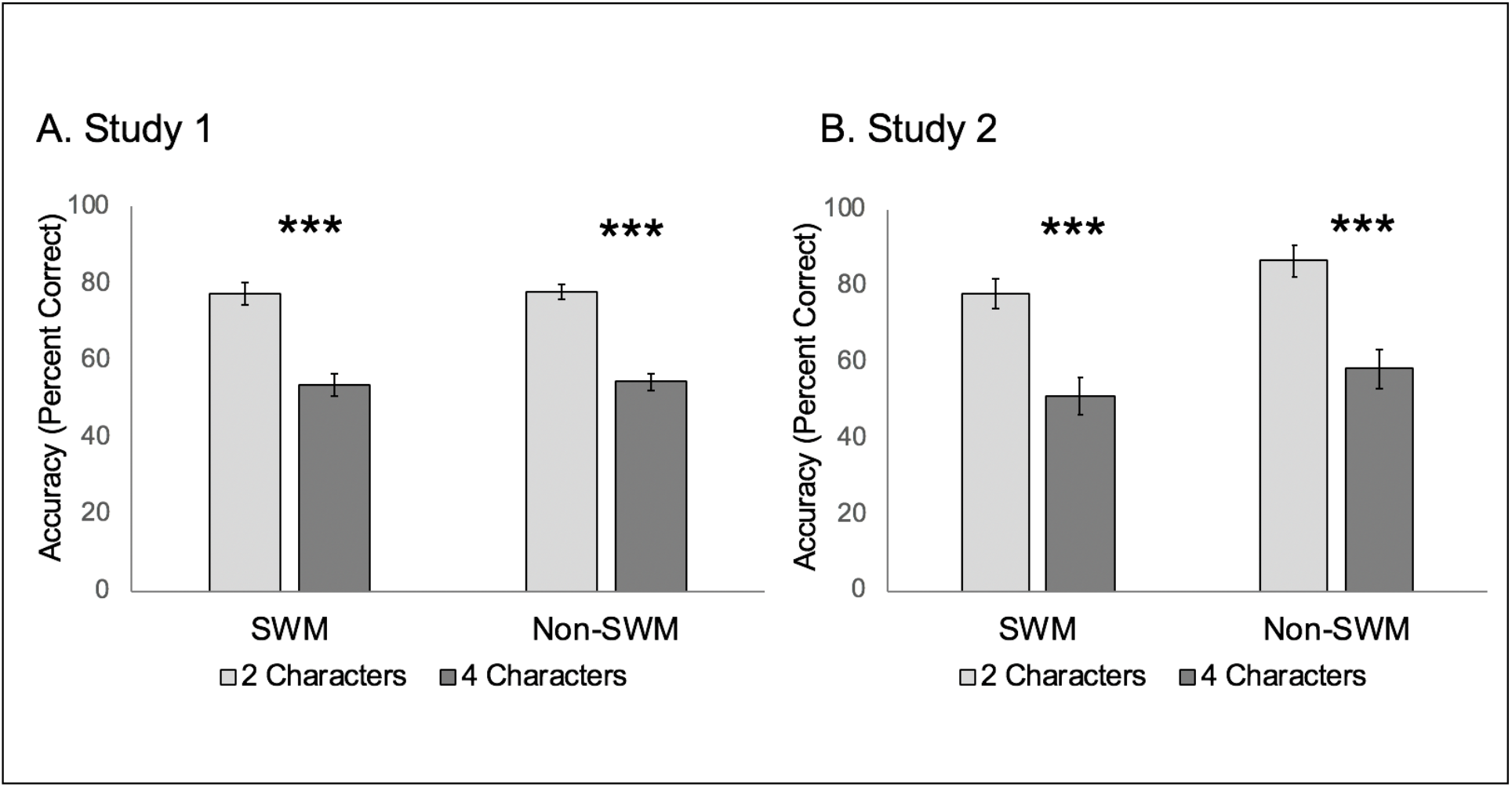
Task accuracy in Study 1 (Panel A) and Study 2 (Panel B). In both studies, accuracy was matched for the SWM and non-SWM trials as a function of load level. Additionally, four-load trials were significantly more challenging than two-load trials. Asterisks (***) indicate p<.001.

Reaction time (RT) was defined as the speed (in seconds) with which participants made correct responses. Although participants’ accuracy was equivalent on the SWM and non-SWM trials, RT varied across conditions. There was a significant content (SWM vs. non-SWM) x load level (two-load vs. four-load) interaction (*F*(1,36)=31.21, *p*<.0001, np2=.48), such that the decrement in speed for the four-load vs. two-load trials was larger for non-SWM trials (two-load M=.98, SD=.26; four-load M=1.74, SD=.36) than SWM trials (two-load M=.88, SD=.32; four-load M=1.11, SD=.65). There was also a main effect of content (SWM vs. non-SWM, *F*(1,36)=28.56, *p*<.0001, np2=.46), such that participants responded more quickly to SWM trials (M=1.00, SD=.47) than non-SWM trials (M=1.34, SD=.24). As is the case for task accuracy, there was also a main effect of load level (two-load vs. four-load, *F*(1,36)=131.32, p<.0001, np2=.79) such that participants responded more quickly to two-load (M=.93, SD=.25) than four-load trials (M=1.40, SD=.41). RT did not statistically differ between subjects who had previously seen the television show *Orange is the New Black* compared to those who had not (*p*’s across load level comparisons>.481; Cohen’s d’s<.26).

#### Whole-Brain Results

The primary goal of the neuroimaging analyses was to test whether default network regions are differentially associated with SWM and non-SWM load. To test this, we searched for clusters of activity more strongly associated with four-load (vs. two-load) SWM trials relative to four-load (vs. two-load) non-SWM trials. This analysis revealed clusters in the dorsomedial subsystem of the default network, including DMPFC, TPJ, MTG extending into TP, and IFG; Figure 3A, Table 1). A cluster outside of the default network in the putamen also emerged in this analysis. The reverse contrast (greater activity for four-load (vs. two-load) non-SWM trials relative to four-load (vs. two-load) SWM trials) showed clusters of activation in supplementary motor area and precentral gyrus (Table 1).

**Figure 3.**
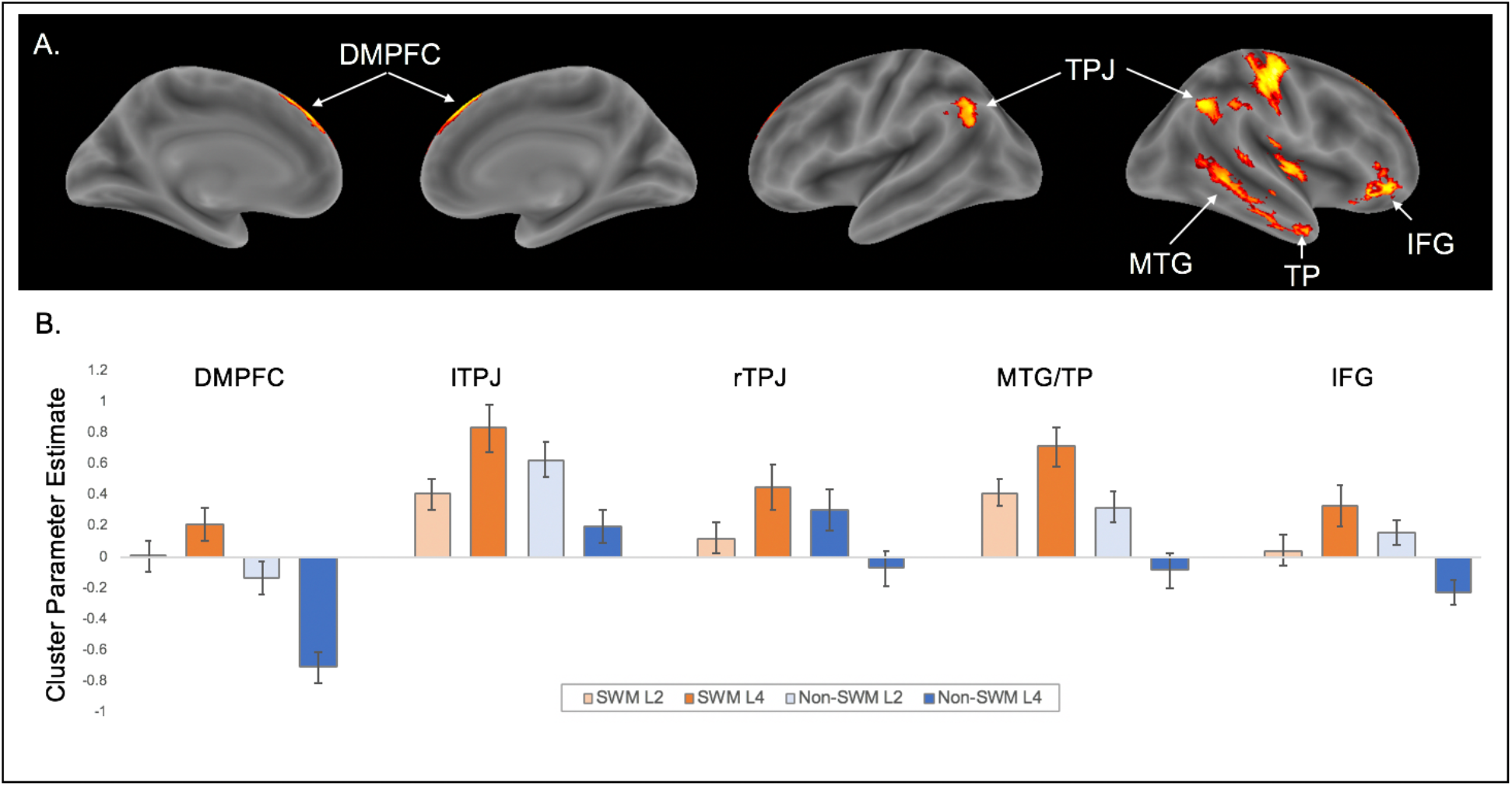
Neuroimaging results for the interaction contrast testing for activity associated with four-load (vs. two-load) SWM trials relative to four-load (vs. two-load) non-SWM trials. Panel A shows clusters in the dorsomedial subsystem of the default network (DMPFC, TPJ, MTG, TP, IFG). Panel B shows parameter estimates from each of these clusters for each condition.

**Table 1.**
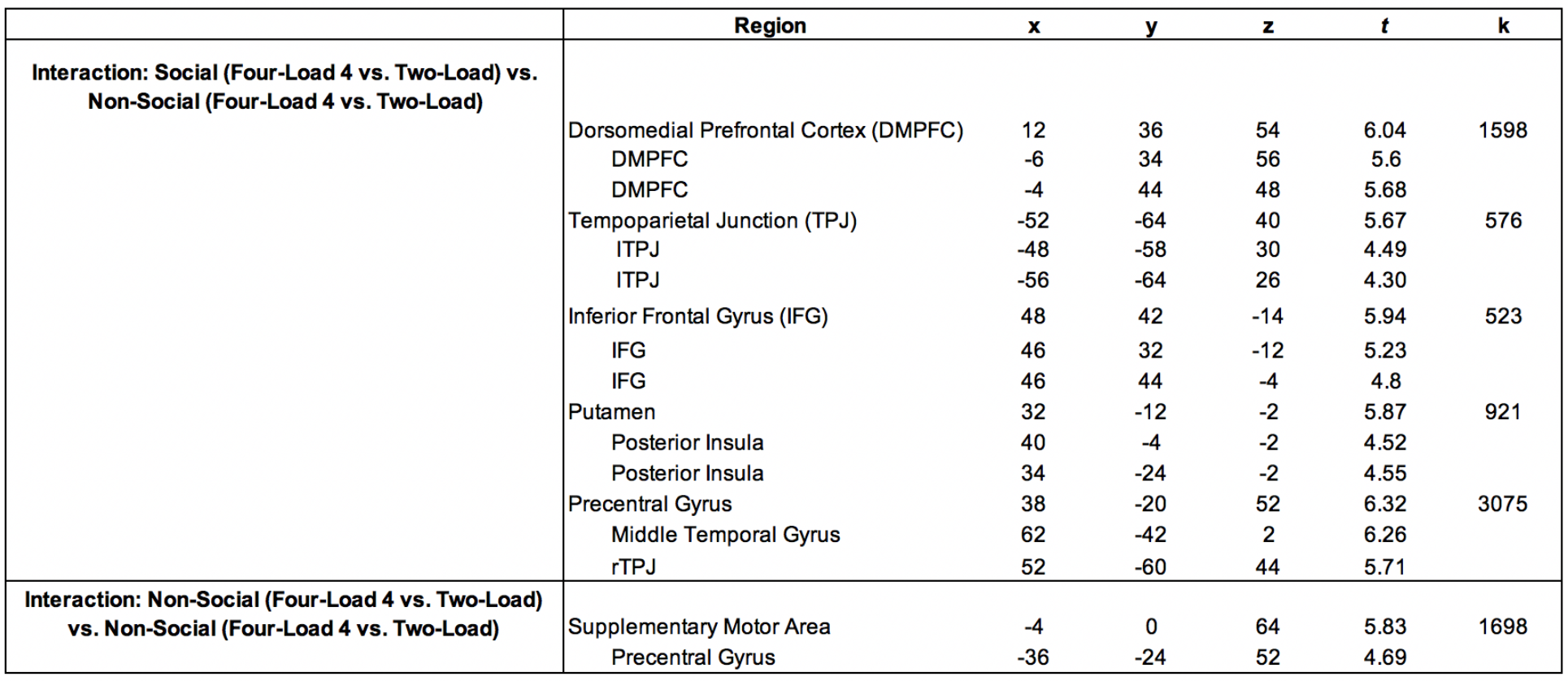
Clusters of neural activity more strongly associated with four-load (vs. two-load) SWM trials relative to four-load (vs. two-load) non-SWM trials (top) and four-load (vs. two-load) non-SWM trials relative to four-load (vs. two-load) SWM trials (bottom). Columns x, y, and z refer to cluster coordinates, t refers to the t-statistic associated with the cluster, and k refers to the cluster extent (number of voxels).

|  | Region | x | y | z | t | k |
| --- | --- | --- | --- | --- | --- | --- |
| **Interaction: Social (Four-Load 4 vs. Two-Load) vs. Non-Social (Four-Load 4 vs. Two-Load)** | Dorsomedial Prefrontal Cortex (DMPFC) | 12 | 36 | 54 | 6.04 | 1598 |
|  | DMPFC | -6 | 34 | 56 | 5.6 |  |
|  | DMPFC | -4 | 44 | 48 | 5.68 |  |
|  | Tempoparietal Junction (TPJ) | -52 | -64 | 40 | 5.67 | 576 |
|  | ITPJ | -48 | -58 | 30 | 4.49 |  |
|  | ITPJ | -56 | -64 | 26 | 4.30 |  |
|  | Inferior Frontal Gyrus (IFG) | 48 | 42 | -14 | 5.94 | 523 |
|  | IFG | 46 | 32 | -12 | 5.23 |  |
|  | IFG | 46 | 44 | -4 | 4.8 |  |
|  | Putamen | 32 | -12 | -2 | 5.87 | 921 |
|  | Posterior Insula | 40 | -4 | -2 | 4.52 |  |
|  | Posterior Insula | 34 | -24 | -2 | 4.55 |  |
|  | Precentral Gyrus | 38 | -20 | 52 | 6.32 | 3075 |
|  | Middle Temporal Gyrus | 62 | -42 | 2 | 6.26 |  |
|  | rTPJ | 52 | -60 | 44 | 5.71 |  |
| **Interaction: Non-Social (Four-Load 4 vs. Two-Load) vs. Non-Social (Four-Load 4 vs. Two-Load)** | Supplementary Motor Area | -4 | 0 | 64 | 5.83 | 1698 |
|  | Precentral Gyrus | -36 | -24 | 52 | 4.69 |  |

Collapsing across load level, we found that multiple regions in the default network (DMPFC, VMPFC, left PC/PCC, lTPJ, and bilateral TP) showed greater activity in response to the SWM vs. non-SWM trials (Supplementary Figure 2, Supplementary Table 2). These results further support the idea that the default network is associated with SWM, and mental state inference more generally. Collapsing across working memory content, and replicating past working memory research, clusters in DLPFC and the precuneus extending into bilateral inferior parietal lobule, showed greater activity in response to four-load vs. two-load trials (Supplementary Figure 2, Supplementary Table 2). For the interested reader, clusters that emerged in the comparison of non-SWM vs. SWM included supplementary motor area, anterior insula, palladium, precuneus, and the cerebellum (Supplementary Figure 2, Supplementary Table 2). Only MPFC and superior temporal lobe showed relatively greater activity to two-load vs. four-load trials (Supplementary Figure 2, Supplementary Table 2). Follow-up analyses showed that all clusters that differentiated SWM and non-SWM trials remained significant when analyses included, for each condition, RT as a parametric modulator (Supplementary Figure 3), suggesting observed differences in neural activity cannot be explained by differences in RT between SWM and non-SWM trials.

#### Pre-defined Default Network Subsystem Results

Next, we assessed whether our whole-brain results replicate in a pre-defined set of default network subsystems. Specifically, we used the default network subsystems identified by Yeo et al. (2011), which includes dissociable regions for the dorsomedial subsystem, core subsystem, and MTL subsystem (Figure 4A). We again observed a significant content (SWM vs. non-SWM) x load level (two-load vs. four-load) interaction in the default network’s dorsomedial subsystem (*F*(1, 36)=11.356, *p*=.002, np2=.240). Follow-up paired samples t-tests confirmed that the dorsomedial network was significantly more active during SWM four-load vs. SWM two-load trials (*t*(36)=2.273, *p*=.029). The dorsomedial subsystem was also significantly less active during non-SWM four-load vs. non-SWM two-load trials (*t*(36)=2.68, *p*=.011). There was also a main effect of content (*F*(1,36)=4.907, *p*=.033, np2=.120) such that the dorsomedial subsystem was relatively more active during SWM (*M*=.29, *SD*=.24) vs. non-SWM trials (*M*=.11, *SD*=.33). There was no main effect of load level (*F*(1, 36)=.281, *p*=.599, np2=.008).

**Figure 4.**
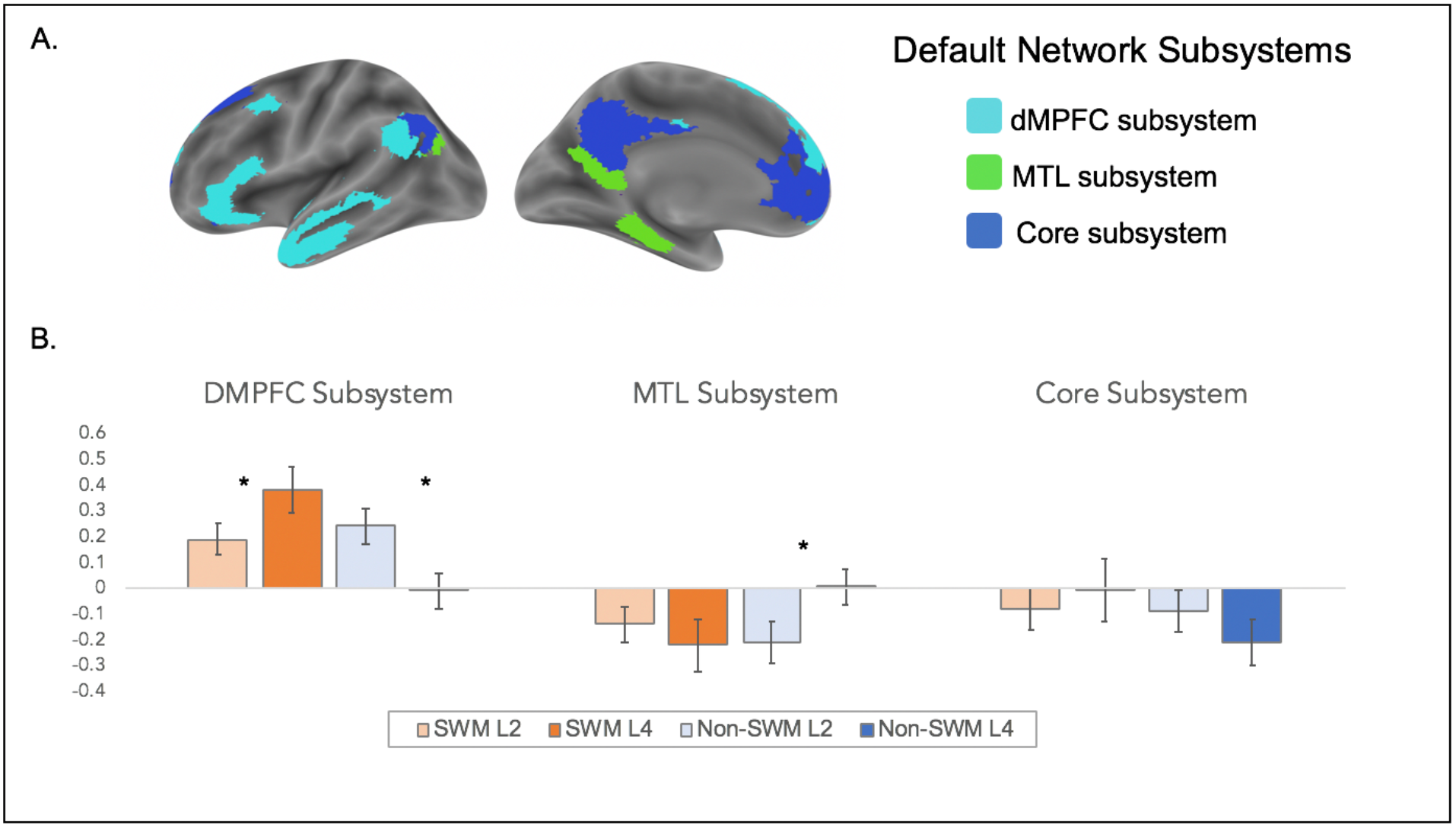
Results from the default network subsystem analyses. Panel A shows the default subsystems, which were generated from the resting state data of 1,000 participants (Yeo et al., 2011). Panel B shows activation in the default subsystems across the social and non-social working memory trial types.

The core subsystem did not show a significant interaction *F*(1, 36)=1.997, *p*=.166, np2=.053), nor a main effect of content (*F*(1, 36)=1.682, *p*=.203, np2=.045) or load level (*F*(1,36)=.202, *p*=.656, np2=.006). Although the MTL subsystem showed a significant content x load level interaction (*F*(1, 36)=5.578, *p*=.024, np2=.134), this effect was driven by differences in activity in the non-SWM four-load vs. two-load trials (*t*(36)=2.110, *p*=.042), but not SWM four-load vs. two-load trials (*t*(36)=0.802, *p*=.427). The MTL subsystem also did not show a main effect of content (*F*(1, 36)=1.070, *p*=.308, np2=.029) or load level (*F*(1,36)=.601, p=.443, np2=.016). Subsystem results are displayed in Figure 4B.

The goal of Study 1 was to test whether default network regions show relative increases in activity as a function of the number of mental states managed in working memory. The dorsomedial subsystem of the default network showed relative gains in activation as a function of the number of mental states managed in working memory, despite showing relative decreases in activity as a function of the number of names alphabetized in working memory. Critically, SWM and non-SWM trials were matched on task accuracy and these neural patterns persisted when controlling for differences in RT across conditions. Collectively, these results suggest that engaging default network regions during SWM generalizes to managing mental state inferences in working memory. Because our task measures SWM performance objectively and given past suggestions that SWM skill may relate to social success (Meyer et al., 2012; Meyer et al., 2015; Krol et al., 2018; Smith et al., 2018; Mikels et al., 2019), our next goal addressed in Study 2 was to assess whether individual differences in SWM preferentially relate to individual differences in social integration.

## Study 2

### Participants

Eighty-eight participants (Males=44, Females=44; average age=23.53 years, SD=4.49,66% Caucasian, 16% Asian, 9% Black or African American, 7% Hispanic or Latino, 2% Other) completed Study 2. Given research showing that social network size decreases during the years following young adulthood (English & Carstensen, 2014), to avoid age confounds in our results we constrained recruitment to participants between the ages of 18-29. Sample size was determined by a power analysis in G*Power 3.1 (Faul et al., 2007), which showed that at least 85 participants were needed to allow for 80% power to detect a medium effect size (i.e. Pearson r = .30) when examining the correlation between task performance and social network size. Participants completed the study for course credit or monetary payment ($6).

### Procedure

#### Social Working Memory Paradigm

Participants completed the SWM paradigm developed for Study 1. Participants watched the *Orange is the New Black* montage and indicated whether or not they had seen the television show in the past. Next, they received task instructions, completed practice trials, and finally performed the experimental task consisting of 36 randomly presented trials (9 per condition).

#### Social Integration

To assess social integration, we used a questionnaire developed by Stiller & Dunbar (2007) which measures the number of social network members interacted with over the past seven days. In this measure, participants are asked to list all of the people with whom they had personal contact or communication within the past 7 days, excluding i) work colleagues seen only in a work environment (unless they considered them genuine friends), ii) contacts with professionals for appointments (such as doctors), and iii) other casual acquaintances (e.g., brief encounters in a shop).

### Results

#### Task Performance

We replicated the task accuracy findings observed in Study 1. There was neither a significant content (SWM vs. non-SWM) x load level (two-load vs. four-load) interaction (*F*(1, 87)=1.67, *p*=.200, np2=.02), nor a significant main effect of content (SWM vs. non-SWM; *F*(1, 87)=1.36, p=.248, np2=.02; Figure 2B). These findings again suggest that the SWM and non-SWM trials are matched on accuracy. There was a significant main effect of load level (two-load vs. four-load; *F*(1,87)=139.32, *p*<.0001, np2=.61) such that four-load trials (M=75%, SD= 19%) were more challenging than two-load trials (M=54%, SD=20%). As in Study 1, task accuracy did not statistically differ between participants who had previously seen *Orange is the New Black* (N=32) relative to participants who had not (N=39; *p*’s>.407, Cohen’s d’s<.19; *Supplementary Table 1*). Collectively, these results again suggest that difficulty level (in terms of accuracy) was equivalent for the SWM and non-SWM trials, that the four-load manipulation successfully taxes working memory resources, and that previous exposure to the television show *Orange is the New Black* does not moderate task performance.

We also replicated the RT results observed in Study 1. There was a significant content (SWM vs. non-SWM) x load level (two-load vs. four-load) interaction (*F*(1,84)=48.28, *p*<.0001, np2=.37), such that the decrement in speed for the four-load vs. two-load trials was larger for non-SWM trials (two-load M=1.04, SD=.41; four load M=1.67, SD=.37) than SWM trials (two-load M=.73, SD=.29; four-load M=.92, SD=.39). There was a main effect of content (SWM vs. non-SWM, *F*(1, 84)=255.47, *p*<.0001, np2=.75), such that participants responded more quickly to SWM (M=.82, SD=.29) than non-SWM trials (M=1.35, SD=.26). As is the case for task accuracy, there was also a main effect of load level (two-load vs. four-load, *F*(1, 84)=185.48, *p*<.0001, np2=.69) such that participants responded more quickly to two-load (M=.89, SD=.24) vs. four-load trials (M=1.30, SD=.29). RT did not statistically differ between subjects who had previously seen *Orange is the New Black* compared to those who had not (*p*’s across load level comparisons>.105; Cohen’s d’s<.40).

#### Relationship Between SWM and Social Integration

Interaction with social network members significantly correlated with SWM accuracy on the two-load trials (*r*=.21, *p*=.045) and four-load trials (*r*=.31, *p*=.003, Figure 5A). A regression analysis demonstrated that the relationship between social network member interaction and four-load SWM accuracy remained significant when controlling (i.e., adding as a covariate) two-load SWM accuracy (SWM L4 ***β***=.30, *t*=2.23, *p*=.028; SWM L2 ***β***=.03, *t*=.21, *p*=.834). Thus, SWM capacity, above and beyond mental state inference in general, is associated with social network integration. In contrast, interaction with social network members did not significantly correlate with non-SWM accuracy on the two-load (*r*=.05, *p*=.643, Supplementary Figure 4) or four-load trials (*r*=.17, *p*=.108, Supplementary Figure 4). That said, it is noteworthy that the relationship between social integration and non-SWM four-load trials, which can be considered the non-SWM ‘yoked control condition’ for the SWM four-load trials, showed a trend in the positive direction. We therefore ran a follow-up regression analysis to investigate whether the relationship between social network integration and SWM four-load accuracy persists when controlling for non-SWM four-load accuracy. This was indeed the case (four-load SWM accuracy ***β***=.28, *t*=2.62, *p*=.010) and is consistent with the primary results showing that SWM is associated with social integration.

**Figure 5.**
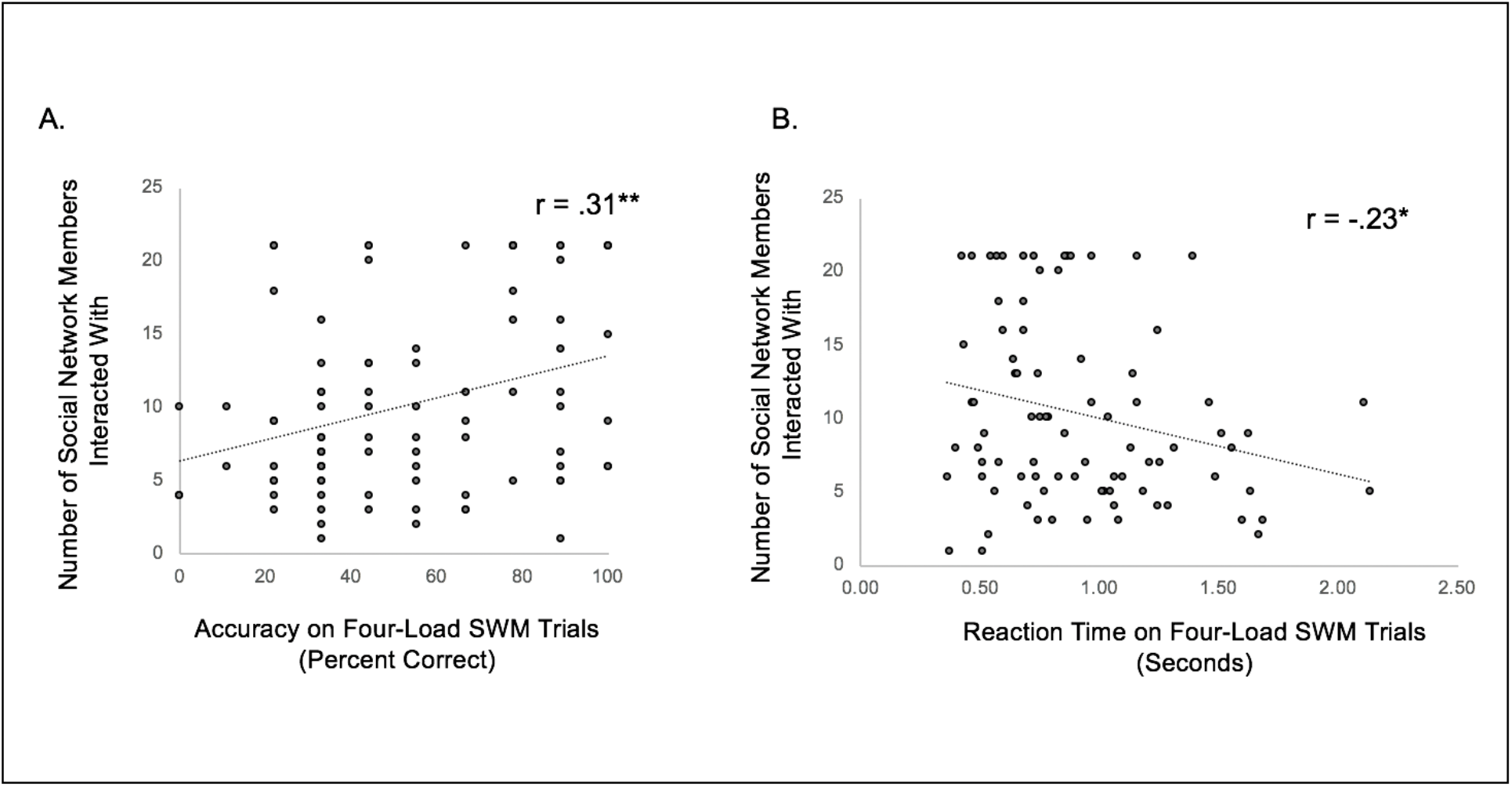
Relationship between social working memory (SWM) task performance on four-load trials and the number of social network members interacted with over a seven-day period. Panel A shows results for task accuracy and Panel B shows results for reaction time (RT). ** indicates p<.005 and * indicates p<.05.

Reaction time analyses paralleled those observed for task accuracy. Individuals who interacted with the most social network members were also faster to make a correct response to SWM four-load trials (*r*=-.23, *p*=.032, Figure 5B) and marginally SWM two-load trials (*r*=-.19, *p*=.073). In contrast, interaction with social network members was not significantly related to reaction time on non-SWM four-load trials (*r*=-.09, *p*=.393, Supplementary Figure 4) nor two-load trials (*r*=-.17, *p*=.107, Supplementary Figure 4).

## Discussion

How do we manage mental state inferences in working memory? Here we found that the dorsomedial subsystem of the default network shows relative increases in activity as a function of the number of mental states considered in working memory. In contrast, these regions showed relative decreases in activity as a function of non-mental state information (the number of names alphabetized) considered in working memory. Such findings are consistent with the suggestion that the default network is differentially associated with SWM and non-SWM (Meyer et al., 2012; 2015). Moreover, only individual differences in SWM task performance, specifically on trials that posed the greatest SWM load, correlated with social integration. Thus, SWM capacity, above and beyond mental state inference and non-SWM capacity more generally, may help us navigate everyday social life.

Although the default network comprises multiple subsystems, here we found that only the dorsomedial subsystem showed relative increases in activity with the number of mental states managed in working memory. These findings are consistent with past research on mental state inference, which consistently implicate TPJ and DMPFC, key nodes in the dorsomedial subsystem, in assessing others’ minds (Dodell-Feder, Koster-Hale, Bedny, & Saxe, 2011; Gallagher, Happe, Brunswick, Fletcher, Frith, & Frith, 2000; Schurz, Radua, Aichhorn, Richlan, & Perner, 2014). Relevant to understanding the role of different default network regions in SWM, graph analytic methods show that the default network is comprised of three subsystems: the core subsystem comprising MPFC, PC/PCC, and PIPL, the medial temporal lobe subsystem comprising hippocampal formation, retrosplenial cortex and dPIPL, and the dorsomedial subsystem comprising DMPFC, TPJ, and MTG extending into TPs (Yeo et al., 2011). Thus, it is possible that different default subsystems are preferentially associated with maintaining and manipulating different types of social information in working memory.

For example, MPFC and PC/PCC are consistently implicated in self-referential processing, as well as thinking about close others, such as friends and family (Denny, Kober, Wager, & Ochsner, 2012; Krienen, Tu, & Buckner, 2010). Previous SWM research found that MPFC increased activity as a function of the number of friends’ traits considered in working memory (Meyer et al., 2012; 2015). In contrast, here we observed that MPFC was less active in response to the higher (vs. lower) social and non-social load trials (i.e., the comparison of four-load vs. two-load trials, collapsed across SWM and non-SWM trials). Our findings suggest that MPFC disengages in response to working memory load when managing social information about *strangers*, unlike when managing social information about *close others*. To further clarify the role of the default network subsystems in working memory, it may be helpful for future research to parameterize working memory load across self, close other, and stranger targets. Interestingly, although some research has implicated MPFC in managing one’s own emotional responses to affective stimuli in working memory (Smith, Lane, Sanova, Alkozei, Smith, & Killgore, 2018b; Smith, Lane, Sanova, Alkozei, Bao, Smith, Sanova, Nettles, & Killgore, 2018c; Waugh, Lemus, & Gotlib, 2014), to our knowledge no research has investigated the brain basis of managing self-concept information (e.g., beliefs about one’s personality) in working memory. This gap is surprising given social psychological theory suggesting that managing self-concept information in working memory is important for self-regulation and fulfilling interpersonal relationships (Markus & Wurf, 1986).

The observation that the dorsomedial subsystem preferentially supports mental states in working memory provides new insight into the functional properties of this system and its role in theory-of-mind (TOM), or the capacity to represent others’ mental states. The classic paradigm used to assess TOM is the false-belief test (Wimmer & Perner, 1983; Dufour et al., 2013). In this paradigm, participants first observe person A witness person B place an item in one of two boxes. Person A leaves the room and Person B moves the item to the other box. Person A returns to the room and the participant is asked which box Person A believes the item is in. Although not typically framed as a working memory task, determining a correct answer to false belief tasks requires working memory; participants must maintain Person A’s belief over time to answer correctly. Thus, an alternative interpretation of TPJ and DMPFC activity in response to false-belief tasks is that these regions support the momentary maintenance of others’ mental states in working memory. One impediment to this interpretation from past work is that traditional false-belief tasks do not parameterize working memory load, making it difficult to disentangle whether these regions are important for mental state inference generally versus maintaining mental state inferences in working memory specifically. Because our two-load and four-load SWM trials required mental state inference, but only the four-load trials increased mental state load in working memory, our results provide helpful insight: TPJ and DMPFC show relative increases in activation as a function of the number of mental state demands to working memory, above and beyond mental state inference per se. These findings are consistent with another recent study finding that the TPJ shows greater activity when making two false belief inferences versus one false belief inference (Ozdem, Brass, Schippers, Van der Cruyssen, & Van Overwalle, 2019) and suggest that activity in these regions during false belief tasks may reflect a (social) working memory process.

Our paradigm is also useful for assessing TOM capacity in adults, who traditionally perform at ceiling on most false-belief tasks (Brunet, Sarfati, Hardy-Bayle, & Decety, 2000; Fletcher et al., 1995; Walter, Ciaramidaro, Enrici, Pia, & Bara, 2004). TOM research focuses heavily on when children acquire the capacity to infer mental states (e.g., Wellman, 1992) and how these mechanisms are altered in populations who struggle to understand others, such as individuals with an autism spectrum disorder (ASD; e.g., Baron-Cohen, 2000). However, far less research examines the upper bounds of TOM expertise, perhaps in part because there are little-to-no paradigms that effectively create variability in healthy adult performance. In contrast, participants in Studies 1 and 2 did not perform at ceiling on our tasks, and individual differences in task performance related to real-world social integration. Thus, our paradigm may be an effective way to begin to unpack individual differences in adult TOM and its link to social prowess.

In contrast to our brain imaging findings, other research on managing other people’s mental states in working memory have found increased activity in the lateral frontoparietal network typically associated with working memory (Smith et al., 2017), but relatively suppressed activity in the default network when participants track others’ emotions (Xin & Lei, 2015). Critically, however, this past work uses paradigms in which participants maintain the external facial features of emotional expressions in working memory. For example, participants are asked to indicate whether a given person’s emotional expression matches the emotional expression of the person shown two trials prior (Xin & Lei, 2015). In these paradigms, participants do not need to engage in high-level social inference; maintaining visual features of facial expression is sufficient. In our paradigm, the facial expressions of the individuals in the photographs are held constant from trial-to-trial, but participants must make and manage mental state inferences in working memory. In everyday life, these two processes often unfold in tandem (e.g., observe a facial expression and make a high-level social inference). To gain a more wholistic understanding of social working memory processes in everyday life, an interesting direction for future work may be to assess how external (e.g. facial expressions) and internal (e.g. social inferences) sources of information are integrated in working memory.

The behavioral results from Study 2 complement and extend past paradigms used to assess whether SWM skills relate to social network integration. Past work has shown that the ability to manage people’s personality traits in working memory (Krol et al., 2018) and maintain information about characters’ mental states from a story (Stiller et al., 2007) correlate with the number of social network members interacted with over the course of a week. The proposed interpretation of these results is that skill in maintaining and manipulating people’s internal characteristics in working memory may help us cultivate social networks. However, two limitations impede this interpretation. The first study used the SWM task in which participants rank a set of friends along trait dimensions ‘from most-to-least’ in working memory and determine accuracy by comparing participants’ answers on a given SWM trial to their ratings of their friends’ traits provided two weeks earlier. Thus, task accuracy is computed with subjective assessments of friends’ traits, and variables such as whether a new event changed a participant’s perception of their friend(s) confound the results. The finding that the ability to remember characters’ mental states from a story correlates with social network integration confounds other variables with task performance, most notably reading comprehension. The paradigm developed here overcomes these limitations, as task accuracy is computed objectively and does not rely on reading comprehension (all stimuli are presented visually). Moreover, we found that task accuracy specifically on the four-load SWM trials, above and beyond task accuracy on the two-load SWM trials, uniquely predicts the number of social network members interacted with over a week. This finding further suggests that the ability to maintain and manipulate mental states in working memory, above and beyond mental state inference per se, is associated with greater social network integration. Future work may reveal the causal direction of this relationship, assessing whether SWM skills prospectively predict social network size, whether cultivating larger social networks builds SWM capacity, or both.

### Limitations

While our results provide helpful insight into the brain basis of SWM and its connection to social integration, they are not without limitations. First, although we were successful in developing a paradigm that computes SWM task accuracy more objectively than past tasks, it is noteworthy that our computation of accuracy assumes that all relationship partners’ mental states equally impact one another. However, in real life, different relationship partners may impact another person’s mental state to different degrees. For example, perhaps your romantic partner’s mental states have a larger impact than your friend’s mental states in determining your own. An interesting direction for future research will be to model the varying weights of emotional contingencies between people to more naturalistically assess how the social brain makes such inferences.

Second, although the SWM and non-SWM trials differ in mental state inference, they also differ in the extent to which they rely on new, episodic information acquired from watching the *Orange is the New Black* video montage. Given that the MTL default network subsystem associated with episodic memory (Squire & Zola-Morgan, 1991) did not distinguish SWM from non-SWM trials, we suspect this difference is not driving our results. Nonetheless, future work may test this possibility more directly, for example by comparing SWM responses to managing new vs. old social information. Related to SWM and non-SWM task differences, these trials also differed in reaction time, suggesting the conditions—despite being well-matched on accuracy— may still vary in difficulty. On the one hand, this difference in RT could suggest that the greater default network activity associated with SWM reflects this network’s involvement in less vs. more effortful cognitive processing. On the other hand, this alternative explanation is unlikely, given that our findings persisted even when we controlled for reaction time on a trial-by-trial basis.

Third, our measure of social network integration relies on self-reported social interaction. Although this measure is commonly used in social cognition research (Heleven, & Van Overwalle, 2016; Heleven & Van Overwalle, 2019a; Heleven & Van Overwalle, 2019b; Lewis, Rezaie, Brown, Roberts & Dunbar, 2011; Roberts, Wilson, Fedurek, & Dunbar, 2008), it is imperfect. For example, a person might not remember all of their social interaction partners when probed one week later. Moving forward, future research may replicate and extend these results with other assessments of social network integration, such as those used in formal social network analyses (Parkinson, Kleinbaum, & Wheatley, 2017).

### Conclusion

In summary, we examined how people manage mental state inferences in working memory and whether skill in this domain relates to real-world social integration. Study 1 demonstrated that the dorsomedial subsystem of the default network shows relative increases in activity as a function of the number of mental states considered in working memory, despite relative decreases in activity as a function of the number of non-mental state information considered in working memory. Study 2 showed that skill in managing more mental states in working memory preferentially correlates with social integration. These findings add further support to the idea that SWM relies on partially distinct brain systems and may be critical to interpersonal success.

## Acknowledgments

We would like to thank Joseph Mastromonaco for creating the *Orange is the New Black* montage and Matthew Lieberman for his suggestions on the experimental design. We would also like to thank Thuy-Vy Nguyen and Remy Borinsky for their assistance with data collection and analysis for Study 2.

