## SupplementalMaterials for "Theory of Mind*s*: Managing mental state inferences in working memory is associated with the dorsomedial subsystem of the default network and social integration"

### Supplementary Materials

| | | Not Seen TV Show | | Seen TV Show | | | | |
|---|---|---|---|---|---|---|---|---|
| **Study 1** | | **Mean** | **SD** | **Mean** | **SD** | ***t*** | ***p*** | **Cohen's d** |
| **Accuracy (Percent Correct)** | **SWM Load 2** | 76 | 24 | 81 | 21 | 0.74 | 0.466 | 0.22 |
| | **SWM Load 4** | 46 | 26 | 59 | 35 | 1.37 | 0.178 | 0.42 |
| | **Non-SWM L2** | 85 | 23 | 89 | 23 | 0.49 | 0.629 | 0.17 |
| | **Non-SWM L4** | 55 | 32 | 63 | 22 | 0.79 | 0.436 | 0.29 |
| **Reaction Time (RT)** | **SWM Load 2** | 0.88 | 0.33 | 0.88 | 0.33 | 0.11 | 0.912 | 0.00 |
| | **SWM Load 4** | 1.13 | 0.69 | 1.09 | 0.61 | 0.16 | 0.875 | 0.06 |
| | **Non-SWM L2** | 1.00 | 0.32 | 0.96 | 0.14 | 0.43 | .670 | 0.16 |
| | **Non-SWM L4** | 1.78 | 0.42 | 1.69 | 0.25 | 0.71 | 0.482 | 0.26 |
| **Study 2** | | | | | | | | |
| **Accuracy (Percent Correct)** | **SWM Load 2** | 70 | 27 | 75 | 26 | 0.82 | 0.415 | 0.19 |
| | **SWM Load 4** | 50 | 29 | 58 | 26 | 0.83 | 0.408 | 0.29 |
| | **Non-SWM L2** | 79 | 15 | 79 | 20 | 0.15 | 0.884 | 0.00 |
| | **Non-SWM L4** | 55 | 21 | 58 | 23 | 0.31 | 0.615 | 0.14 |
| **Reaction Time (RT)** | **SWM Load 2** | 0.71 | .20 | 0.71 | 0.31 | 0.01 | 0.994 | 0.00 |
| | **SWM Load 4** | 0.96 | 0.42 | 0.83 | 0.33 | 1.41 | 0.162 | 0.34 |
| | **Non-SWM L2** | 1.03 | 0.27 | 0.98 | 0.26 | .80 | 0.427 | 0.19 |
| | **Non-SWM L4** | 1.73 | 0.4 | 1.58 | 0.35 | 1.64 | 0.106 | 0.4 |

*Note: Note: In Study 2, 17 participants did not indicate whether or not they had seen the televsion show Orange is the New Black*

*Supplementary Table 1. Task performance in Studies 1 and 2 for participants who had and had not previously seen the television show Orange is the New Black.*

| | Region | x | y | z | t | k |
|---|---|---|---|---|---|---|
| **SWM vs. Non-Social** | | | | | | |
| | Dorsomedial Prefrontal Cortex (DMPFC) | -8 | 54 | 36 | 9.38 | 2335 |
| | DMPFC | 14 | 38 | 54 | 7.23 | |
| | DMPFC | -10 | 58 | 26 | 7.00 | |
| | Ventromedial Prefrontal Cortex (DMPFC) | -2 | 34 | -20 | 7.44 | 468 |
| | VMPFC | 0 | 26 | -22 | 7.32 | |
| | VMPFC | -2 | 50 | -16 | 6.59 | |
| | Precuneus/Posterior Cingulate (PC/PCC) | -2 | -50 | 28 | 8.41 | 1212 |
| | PC/PCC | 0 | -68 | 36 | 4.53 | |
| | left Temporal Pole (lTP) | -50 | 10 | -32 | 7.97 | 1247 |
| | lTP | -62 | -10 | -22 | 6.31 | |
| | lTP | -60 | -2 | -24 | 5.99 | |
| | right Temporal Pole (rTP) | 48 | 12 | -36 | 6.57 | 531 |
| | rTP | 52 | 16 | -28 | 5.34 | |
| | rTP | 62 | -6 | -18 | 4.59 | |
| | left Temporoparietal Junction (lTPJ) | -54 | -66 | 24 | 4.93 | 352 |
| | lTPJ | -42 | -62 | 24 | 4.71 | |
| | lTPJ | -50 | -66 | 16 | 4.37 | |
| **Non-Social vs. Social** | | | | | | |
| | Supplementary Motor Area | -2 | 2 | 64 | 8.57 | 13085 |
| | Superior Frontal Gyrus | 26 | 10 | 54 | 8.42 | |
| | Middle Frontal Gyrus | -26 | 6 | 58 | 8.06 | |
| | Anterior Insula | 36 | 16 | 6 | 6.41 | 720 |
| | Inferior Frontal Gyrus | 34 | 26 | -6 | 3.98 | |
| | Anterior Insula | 32 | 18 | -10 | 3.58 | |
| | Palladium | -10 | 0 | 0 | 5.05 | 2183 |
| | Putamen | -18 | 0 | 14 | 4.93 | |
| | Thalamus | 14 | -8 | 10 | 4.87 | |
| | Precuneus | -10 | -60 | 54 | 7.74 | 5099 |
| | Precuneus | 12 | -62 | 58 | 7.36 | |
| | Precuneus | 4 | -42 | 46 | 6.15 | |
| | Cerebellum | -30 | -52 | -30 | 6.43 | 1139 |
| | Cerebellum | -18 | -54 | -22 | 5.16 | |
| | Cerebellum | -24 | -62 | -24 | 5.14 | |
| | Cerebellum | 30 | -58 | -30 | 6.41 | 426 |
| | Cerebellum | 40 | -56 | -32 | 5.59 | |
| | Cerebellum | 10 | -72 | -22 | 4.11 | |
| **Four-Load vs. Two-Load** | | | | | | |
| | Dorsolateral Prefontal Cortex (DLPFC) | 28 | 0 | 54 | 11.70 | 14363 |
| | Supplementary Motor Area | -6 | 14 | 44 | 10.05 | |
| | Dorsal Anterior Cingulate Gyrus | 6 | 18 | 42 | 9.79 | |
| | Precuneus | 4 | -60 | 52 | 11.13 | 21139 |
| | Supramarginal Gyrus | 44 | -36 | 44 | 10.54 | |
| | Superior Parietal Lobule | -22 | -66 | 50 | 10.51 | |
| | Palladium | -16 | -2 | 2 | 7.11 | 2905 |
| | Thalamus | 14 | -8 | 8 | 6.10 | |
| **Two-load vs. Four-Load** | | | | | | |
| | Medial Prefrontal Cortex (MPFC) | -6 | 52 | -2 | 5.02 | 1901 |
| | MPFC | -2 | 32 | -2 | 5.00 | |
| | MPFC | -10 | 36 | -8 | 5.00 | |
| | Superior Temporal Lobe | -44 | -6 | -8 | 5.18 | 430 |
| | Posterior Insula | -40 | -8 | 18 | 4.42 | |
| | Superior TP | -40 | 0 | -16 | 4.27 | |

*Supplementary Table 2. Clusters of neural activity associated with Social vs. Non-Social working memory trials and Four-load vs. Two-Load working memory trials.*

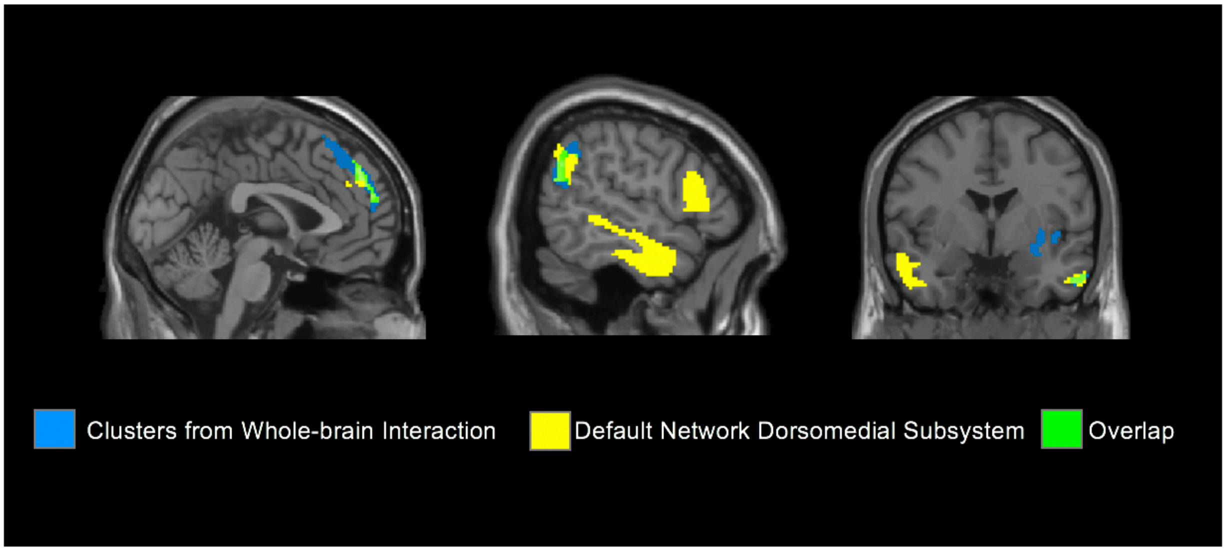


*Supplementary Figure 1. Clusters of neural activity from the whole-brain interaction comparing greater activity in response to SWM four-load (vs. two-load) relative to non-SWM four-load (vs. two-load) trials (in blue), the dorsomedial subsystem of the default network defined by Yeo et al. (2011; yellow) and their overlap (green). There is overlap in dorsomedial prefrontal cortex, tempoparietal junction, and temporal poles.*

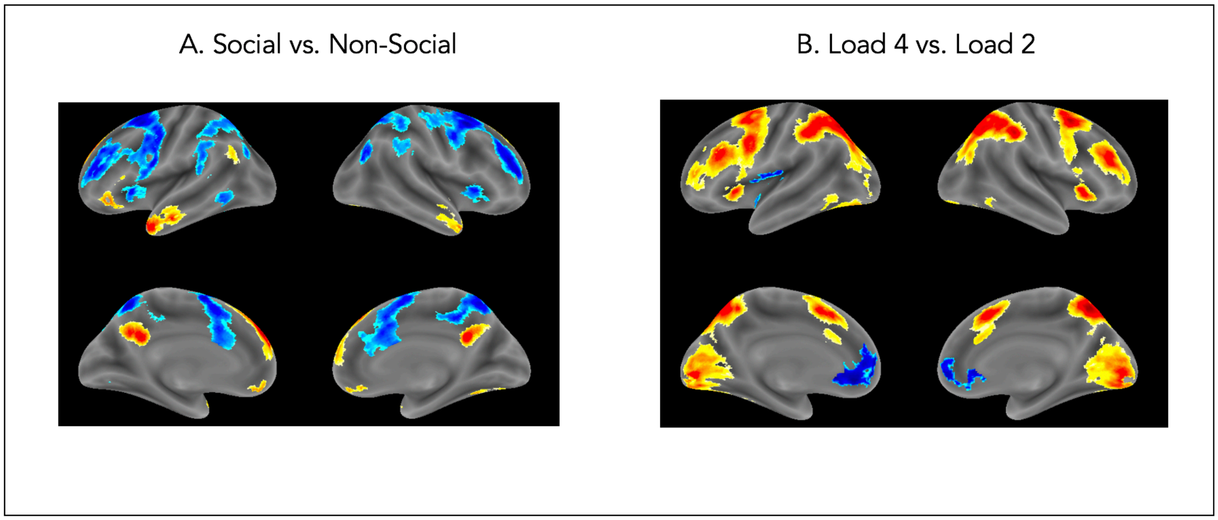


*Supplementary Figure 2. Main effect analyses. Warm colors indicate activation increases and cool colors indicate activation decreases. Panel A shows neural activity associated with SWM vs. non-SWM trials. Panel B shows neural activity associated with four-load vs. two-load trials.*

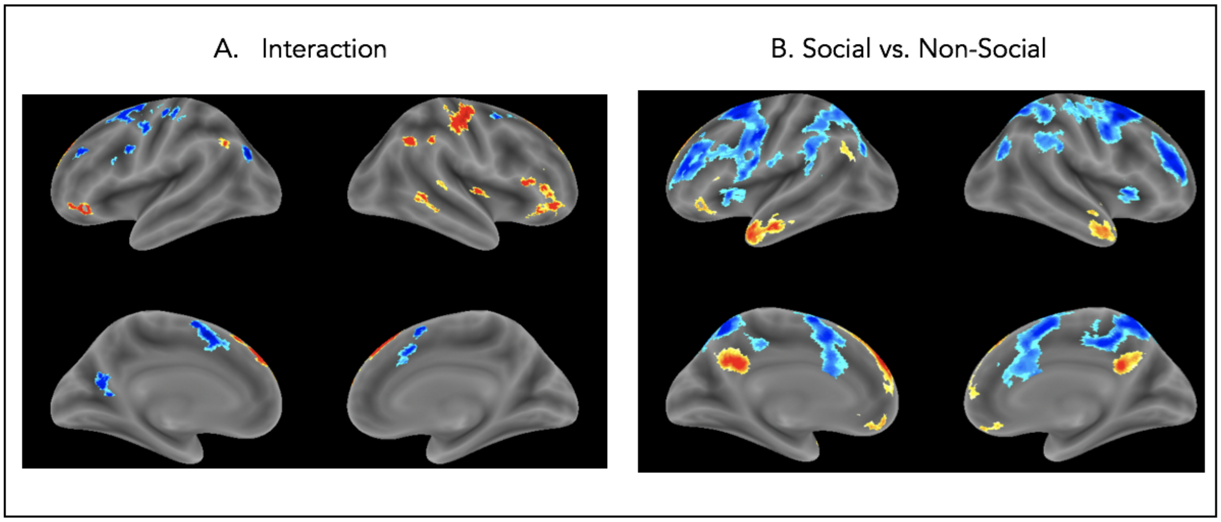


*Supplementary Figure 3. Contrasts in which SWM and non-SWM trials showed different patterns of neural activity, controlling for reaction time (RT). Warm colors indicate activation increases and cool colors indicate activation decreases. Panel A shows results for the interaction contrast testing for neural activity more strongly associated with SWM four-load (vs. two-load) relative to non-SWM four-load (vs. two-load) trials. Panel B shows results for the contrast comparing SWM relative to non-SWM trials, collapsed across load level.*

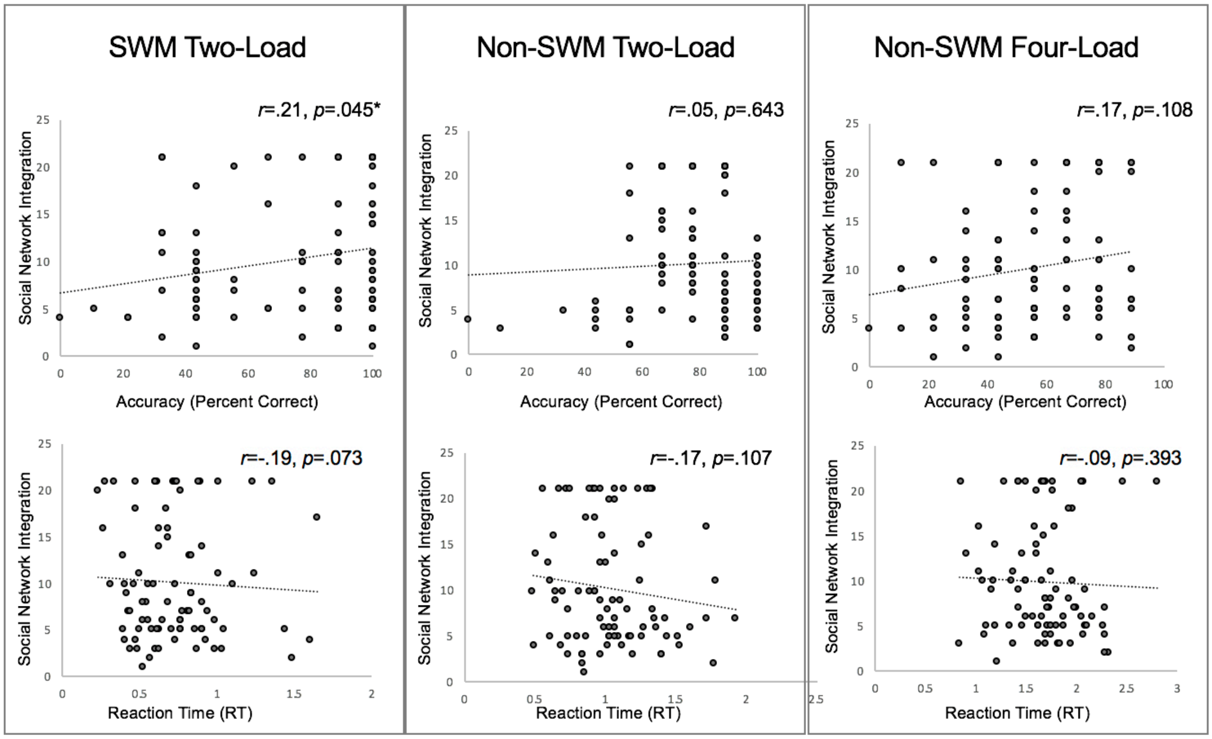


*Supplementary Figure 4. Scatter plots showing the relationship between social network integration (y-axis) and SWM Two-Load, non-SWM Two-Load and non-SWM Four-Load trials.*
